# Eurasian Spoonbill chicks receive parental care up to several months after fledging, but not into migration

**DOI:** 10.1101/2025.05.15.654226

**Authors:** Tamar Lok, Petra de Goeij, Eldar Rakhimberdiev, Theunis Piersma, Wouter Vansteelant

## Abstract

Despite its potential role in affecting survival, habitat use and migration strategies of juvenile birds, post-fledging parental care is poorly studied, as it requires that families can be followed over large distances. Here we combine visual observations of colour-ringed chicks being fed after fledging with GPS-tracking and accelerometer-based behavioural classification of fledged chicks and their parent(s) to quantify post-fledging parental care in Eurasian spoonbills in relation to age of the chick and sex of the chick or parent. We show that the number of observed feedings and the amount of overall contact between chicks and parents, i.e. when chick and parent were <10 m apart, strongly decreased with chick age. Chicks were observed being fed until 125 d old, always within 40 km of the natal colony. The amount of contact strongly varied among the 16 GPS-tagged chick-parent pairs, on average decreasing from 8% of contact time at 40 d old to <2% at 90 d. The last contact occurred at chick ages of 44-136 d (median: 88 d). All contact occurred within 18 km of the natal colony except for the first outbound migratory flight of one chick-parent pair. Both mothers and fathers engaged in post-fledging care, with some evidence that mothers had slightly more contact with chicks than fathers, overall as well as while the chick was begging or foraging. In 10 out of 11 cases that both chick and parent embarked on autumn migration, they did not migrate together: post-fledging care ended on average one week before the adult’s departure and four weeks before the chick’s departure on autumn migration.

**Significance statement:** Prolonged parental care may have positive effects on the survival and influence the habitat use and migration strategies of juvenile birds, as parents may continue to feed their young, show them suitable sites for foraging or resting or suitable routes for migration. Nonetheless, the duration of post-fledging parental care is poorly studied as it requires that parents and their young can be followed over large distances. Based on feeding observations and GPS-tracking of fledged chicks and their parent(s), we show that in Eurasian spoonbills *Platalea leucorodia*, the duration of post-fledging parental care is highly variable but ends before the onset of migration. This has important implications for the development of migration routines of juvenile spoonbills.

## Introduction

Studies on parental care in birds have strongly focussed on the period when birds are bound to the nest (nest building, egg laying, incubation and feeding of chicks), likely due to the difficulty of following families after fledging, when they often travel large distances. After fledging, however, chicks must acquire foraging skills (Mellink et al. 2014), learn where to find suitable sites and habitats for foraging and resting, how to interact with conspecifics and other species, how to avoid predators and, in case of migratory species, acquire suitable migration routines. Not surprisingly, perhaps, young birds often experience considerable mortality shortly after fledging (Naef-Daenzer and Grüebler 2016). Extended parental care during this period has been shown for a variety of species and has likely evolved to reduce this post-fledging mortality by (1) continued feeding of the chicks while they improve their own foraging skills (Mellink et al. 2014), (2) guiding them to suitable sites and habitats for foraging and resting, (3) protecting them from conspecifics, other species and predators and/or (4) guiding them on their first migration, i.e., teaching them suitable migration routines (Chetverikova et al. 2017). An experimental study on barn swallows (*Hirundo rustica*) confirmed that extended post-fledging parental care indeed increased post-fledging survival (Grüebler and Naef‐Daenzer 2010). On the other hand, providing longer post-fledging parental care may go at the expense of future fecundity of the parents (López-Idiáquez et al. 2018), thereby creating a parent-offspring conflict (Rehling et al. 2012; Riou et al. 2012).

While in some species, post-fledging parental care lasts for only a few days, other species show much longer periods of post-fledging parental care, and family associations may even be maintained during and after the first outbound migration (Cramp 1994). While migrating in family groups was long thought to be restricted to Anseriformes (geese, swans; e.g. Scott 1980; Black and Owen 1989; Earnst and Bart 1991), Gruiformes (cranes; e.g. Alonso et al. 2004) and some species of terns (Ashmole and Tovar 1968; Byholm et al. 2022), recent studies revealed that it also occurs in a variety of other species, including great bustard (*Otis tarda*, Palacín et al. 2011), and long-tailed tits (*Aegithalos caudatus*, Chetverikova et al. 2017). In the case of terns, parents also feed their chicks well into the migration, and even as far as the non-breeding grounds (Ashmole and Tovar 1968; Fijn and Kemper 2023). At some point, parents may no longer have a caring role, but family bonds may nonetheless be maintained when the young may help with defending a (breeding or food) territory, warning for potential danger (Black and Owen 1989), or raising genetically related individuals (i.e., cooperative breeding, Koenig and Dickinson 2004). The duration and type of post-fledging parental care (i.e., feeding at nesting site or guiding to suitable sites for foraging and resting) will determine the extent to which juveniles learn suitable habitats and/or migration routes from their own parent(s) or discover these through independent exploration or social interactions with other conspecifics. This will influence the social and genetic structure in a population as well as the capacity of populations to adjust to changing conditions. However, despite its presumed importance, functional role and population-level consequences, only few studies have quantified the duration of post-fledging parental care, as it is difficult to follow families, particularly when they move out of the study area and disperse over large distances (Meyburg et al. 2005).

Here, we investigate post-fledging parental care in Eurasian spoonbills (*Platalea leucorodia leucorodia*). Eurasian spoonbills are colony-breeding birds that show obligate biparental care during egg incubation and chick-rearing prior to fledging (Lok et al. 2024). A recent study in a freshwater population in Hungary showed that after fledging, at an age of *c*. 40 d old, juveniles were still fed by a (presumed) parent at distances up until c. 150 km from the natal colony and at an age of 113 days after hatching, with the last chick feedings observed in early October (Pigniczki 2022). This is considerably longer than what has so far been reported for closely related species (Cramp 1994), which Pigniczki (2022) suggests to be caused by the long time it takes for young spoonbills to master the technique of tactile foraging on often mobile prey. In contrast, the most extreme examples of extended care involve species with rather basic foraging techniques (e.g. grazing geese). As such, the observations from Hungary raise questions about how common it is for spoonbills to extend care for several months after fledging, possibly extending into (part of) the migratory period. If common, such care would have important implications for the development of migratory routines of juvenile birds (Whiten 2019). Spoonbills are known to form post-breeding congregations to prepare for migration (de Boer et al. 2024; Henriques et al. 2025) and to migrate in often mixed-age flocks (Lagarde and Piersma 2021). However, it remains to be confirmed whether (and how often) parents and young migrate together in these flocks, and hence, whether juveniles might learn routes, stop-overs and migratory destinations from their parents.

We combine GPS-tracking and acceleration-based behavioural classification of parents and their young with visual observations of feedings of colour-ringed chicks (of known age and origin) to quantify until what age and what distance from the colony spoonbill chicks receive post-fledging parental care (in the form of being fed or being guided to post-breeding sites or on migration), and whether this depends on the sex of the chick or the parent. As males are the larger sex in spoonbills, male chicks require more food during growth (Krijgsveld et al. 1998) and may therefore be more dependent on still receiving food from their parents after fledging compared to female chicks. In addition, fathers and mothers may differ in the amount of post-fledging parental care they provide (Palacín et al. 2011; Ersoy et al. 2021; Byholm et al. 2022), which could be related to the amount of energy already spent on parental care before fledging, or sex-specific parental care roles that may have evolved.

## Methods

### Study population

For this study, we use observations of colour-ringed Eurasian Spoonbill chicks from colonies throughout The Netherlands between 1999 and 2023 (Photo 1) and GPS and acceleration data of chicks and their parent(s) from the colony on Schiermonnikoog, The Netherlands (53°29□N, 6°15□E) between 2016 and 2018.

### Feeding observations

To retrieve visual observations of feedings of colour-ringed chicks, we selected resightings from the colour-ringing database of the Werkgroep Lepelaar that had feeding-related English or Dutch words in the remarks (i.e., “feed”, “fed”, “voer”) or where feeding was entered as a behaviour. We then further checked the remarks on whether any information was provided about the ring code of the parent (or its chick). This resulted in a dataset of 184 feeding observations of 127 colour-ringed chicks born in The Netherlands between 1999 and 2024, involving 77 colour-ringed parents and 33 feedings by unringed parents.

The sex of a bird was either determined molecularly (see below) or visually. For the birds that were not molecularly sexed, we used the sex identification from visual observations to assess a bird’s (likely) sex. When a bird was visually sexed, it was often noted whether the bird (1) was seen copulating, (2) was seen with partner with clear difference, (3) was seen with partner with unclear difference, (4) was seen alone, (5) has a partner of which sex was molecularly determined, (6) stopped or started incubating at the expected time for a certain sex (starting in the morning or stopping in the evening = male), (7) incubated at night (based on camera trap data) = female. For situations 1, 2, 5 or 7, the identified sex was considered certain, all other visual sex identifications were considered uncertain. For each bird, we then calculated the number of certain and uncertain female and male identifications. If a bird had only certain identifications of a single sex, this was assumed to be the bird’s sex. In all other cases, when the total number of female identifications (certain and uncertain identifications combined) differed by more than two from the total number of male identifications, the most often identified sex was assumed to be the bird’s sex. For 48 of the 54 birds (89%) for which sex was determined both visually and molecularly, the sex was correctly identified based on visual observations using the above-described procedure. The age of the chick was calculated from the length of the 8^th^ primary (P8) at colour-ringing, with the Gompertz-curve formula estimated by Lok et al. (2014): age (days) = −ln(–ln(*P8*/247))/0.095 + 19.3.

It was not possible to record data blind with respect to the birds’ sex because our study involved focal (colour-ringed) individuals in the field. Therefore, observers might sometimes have known (or be able to observe) the sex of the adult birds. However, at the time of these observations, the sex of the chicks had not yet been confirmed molecularly, and visual sex identification was not possible as the chicks have not yet reached their adult size.

### GPS tracking

Between 2012 and 2018, 31 adult spoonbills and between 2016 and 2018, 15 juvenile spoonbills were equipped with a UvA-BiTS GPS/ACC tracker (Bouten et al. 2013). The adult spoonbills were caught on the nest during the late egg incubation stage (for more details, see Lok et al. 2024), while the juveniles were caught shortly before they were able to fly. Upon capture, the birds were ringed with a unique combination of colour-rings and a flag and measured. For molecular sex determination (Fridolfsson and Ellegren 1999), a blood sample of 10-80 μl was collected from the brachial vein and stored in 96% alcohol. The GPS-tracker was mounted on the back of the bird using a non-flexible Teflon harness. The tracker including the neoprene pad and Teflon harness (5 g) weighed 31 ± 12 g (mean ± SD) for the adults and 22 ± 0.9 g for the juveniles. With average body mass being 1646 ± 146 g and 1968 ± 162 g for the tagged adult female and male spoonbills, and 1371 ± 82 g and 1595 ± 133 g for the juvenile female and male spoonbills, the trackers contributed 0.9-2.8% to body mass, thus not exceeding the 3% guideline (Phillips et al. 2003). Data of tagged birds used in this paper are shown in Table S1.

We selected juveniles for GPS-tagging that had at least one GPS-tagged parent. To this aim, the chicks received a uniquely numbered temporary band of flexible plastic around each leg when they were between 5 and 12 days old and still attached to their own nest. During this visit, head-bill length was measured to estimate age (Lok et al. 2014). We then recaptured the chicks when they were 30-32 days old and almost able to fly (at c. 35 days old) and equipped them with a GPS tracker. As the chicks still grow somewhat bigger after fledging (they weighed on average 83% of the adult sex-specific body mass), we left a bit more space between the body and the harness than we did for the adults (i.e. *c*. 2 cm instead of 1 cm space between the tracker and the backbone of the bird). Of the 15 juvenile spoonbills that were GPS-tagged, two died prior to departure on migration and another 5 died later during their first year (Table S1). This resulted in a first-year survival rate of 0.53 ± 0.13 (mean ± SE), which is not significantly different from the first-year survival rate estimated from mark-recapture data and averaged over the period 1988-2008 (0.56 ± 0.04; Lok et al. 2013; Z=-0.20, p=0.84).

For this study, the trackers were set to collect a GPS location every 10 minutes along with a 1.6 s sample of 20 Hz acceleration data. When the birds were within the UvA-BiTS antenna network, all data were downloaded. Between 2012 and 2015, birds were equipped with GPS-trackers whose data could only be downloaded via this antenna network. However, from 2016 onward, we used GPS-GSM trackers that were able to download a small part of the GPS samples via the GSM-network when birds were out of range of the antenna network. This allowed us to also track the birds during their southward migration, albeit at low resolution, as trackers were set to transmit three GPS locations collected during the three hours prior to a GSM session that occurred at 6-hour intervals, provided that the tracker was able to connect to a GSM-network.

The day of departure on autumn migration was defined as the first day that the individual travelled more than 100 km in southward direction between the first and last GPS fix of that day and ended up more than 50 km south of the colony. The precise moment of departure was defined as the date and time associated with the GPS fix preceding the first migratory flight, which was defined as a flight (i.e., a bout of consecutive GPS intervals during which the average speed was at least 5 km/h), with a net travel distance of at least 30 km that started on the day of departure or the preceding day. Including the preceding day was necessary to include the few cases where a bird departed on the evening or night before the departure day as defined above.

To identify contact moments between chick-parent pairs, and the behaviour of the chick during these contact moments, we linked the GPS data of the chick and its parent(s) based on the closest GPS timestamps between the two. We defined the chick to be in contact with its parent when the distance between the two was less than 10 meters. We realize that this is a rather arbitrary boundary distance that likely results in an underestimate of the actual amount of contact as the GPS timestamps of the chick and its parent are never really at the same time. Particularly when birds are walking, foraging or flying, the distance between the locations of the chick and parent quickly becomes larger than 10 m. Therefore, we also did the analysis with a contact distance of 50 m.

### Behavioural classification

Lok et al. (2023) developed a random-forest model for the behavioural classification of spoonbill behaviours from accelerometer data. This model accurately classified resting (sitting and standing), flying and searching (F>0.90) and reasonably accurately identified the ingestion of prey (F=0.73) and walking (F=0.65). Drinking and handling prey were poorly classified (F<0.10). However, this model was trained on acceleration data associated with video-annotated behaviour of adult spoonbills. As such, this model did not include the chick-specific, and for this paper highly relevant, behaviour ‘begging’ for food at a parent, where the chick moves its head up and down (also referred to as ‘pumping’) while standing next to the parent or walking after the parent. Eventually, this begging may be followed by the parent feeding the chick, but as this is such a rarely occurring (only a few times per 24 hours) and very short-lasting (of a few seconds only) event compared to the amount of begging (usually lasting several minutes) preceding this feeding event, we did not attempt to include ‘feeding the chick’ or ‘being fed’ into our behavioural assay. To be able to distinguish ‘begging’ from the accelerometer data, we collected video-footage of GPS-tagged chicks and managed to get 110 s of video-material of begging behaviour, though from one chick only (6381). In addition to this video-footage we visually annotated another 337 s of begging data that directly preceded our video-footages, based on the very distinct acceleration signal of begging. We added these video- and visually annotated ‘begging’ data to the annotated data used in Lok et al. (2023) and trained a random-forest model to distinguish the behaviours ‘search’, ‘handle’, ‘ingest’, ‘sit’, ‘stand’ (also including cases where the bird was moving its body, e.g. when preening, drinking or shaking feathers), ‘fly-active’, ‘fly-passive’, ‘walk’, as covered by Lok et al. (2023)), as well as a new behavioural class ‘beg’. As predictor variables in the models, we used the summary statistics described in Lok et al. (2023), calculated over fixed-length segments (of 0.4 s, 0.8 s or 1.6 s) of the 20 Hz acceleration data measured over three axes, along with GPS-speed. As for the current study, we are primarily interested in distinguishing begging and foraging from other behaviours, we calculated the classification performance (using the F-measure as a balanced measure of sensitivity and precision, see Lok et al. 2023) of the pooled behaviours ‘forage’ (including ‘search’, ‘handle’ and ‘ingest’), ‘rest’ (including ‘sit’ and ‘stand’) and ‘fly’ (including ‘fly-active’ and ‘fly-passive’), and the unpooled behaviours ‘walk’ and ‘beg’. We subsequently applied the best-performing model on the chick data. As walking is often confused with foraging (Lok et al. 2023), we assumed a chick was foraging when it was classified to be foraging on the basis of its accelerometer data and GPS-speed and it was located in foraging habitat. All other classifications of foraging were considered to be walking. We used the habitat map of Lok et al. 2024 to determine whether birds were in foraging habitat (i.e. freshwater, brackish or marine habitats) or on land.

### Data selection and analysis

To select the period after fledging, we only used data of chicks that were at least 35 days old. Using the visual feeding observation dataset, we tested whether the number of observed feedings depended on the chick’s age (randomly selecting one observation per known-age chick, N=127), the chick’s sex (randomly selecting one observation per sexed chick, N=86) or the parent’s sex (randomly selecting one observation per sexed parent, N=57) using generalized linear models with a Poisson error distribution. To have enough counts per age class to make robust statistical inferences, we created chick age classes of 10 days, where 40 = 35-44 days old, 50 = 45-54 days old etc. Moreover, using linear models, we tested whether the distance from the colony where a feeding took place depended on chick age, now treating chick age as unbinned continuous variable.

Of the 15 GPS-tagged juvenile spoonbills, 12 had one GPS-tagged parent while 3 chicks had two GPS-tagged parents, resulting in 18 chick-parent pairs. For the analysis of GPS-based contact moments, we only used data where the absolute difference between the nearest timestamp of the chick and that of its parent was less than 10 minutes, or where the parent had already departed while the chick had not. In the latter case, despite a potentially larger time difference than 10 minutes, it was certain that chick and parent were not in contact. Furthermore, to derive reliable estimates of age-specific proportions of contact between a chick and its parent, we only used days with at least 130 linked GPS locations per chick-parent pair (there would be 144 if the GPS locations occurred at exactly 10-minute intervals and no GPS fixes were missed). For modelling behaviour-specific contact, we only used days with at least 100 linked GPS locations with an associated complete (1.6 s of 20 Hz) acceleration sample for the chick. This dataset is smaller as acceleration measurements were sometimes incomplete or missing, which occurred seemingly randomly, likely due to interference with the process of downloading the data. Finally, we only included chick-parent pairs with at least 10 days of such nearly complete joint data and, due to the much lower data resolution for chicks after their departure on autumn migration, only data prior to departure of the chick. As a result, 16 chick-parent pairs with on average 71 days of nearly complete overall contact and 68 days of behaviour-specific contact data were retained for analysis (Table S1). The oldest chick in this dataset that still had contact with a parent was 136 d old.

Using data of chicks between 35 and maximally 136 d old, we tested whether the proportion of time that chicks have contact, overall or while the chick was begging or foraging, with their parent(s), and the distance to the nest during contact, was explained by chick age, chick sex or parent sex. For this, we used generalized linear mixed models with a binomial (for contact probabilities) and Gamma (for distance to the nest, using a log-link) error distribution and correlated random variation in the intercept and chick age slope among chicks. To avoid overfitting, and as we didn’t have clear expectations about this, we did not consider the two- and three-way interactions between chick age, chick sex and parent sex. While we had not anticipated this beforehand, exploratory analysis revealed strong annual variation in the amount of contact between chick and parent, potentially caused by environmental variation (e.g., food availability). Therefore, we account for this year effect by including it as a fixed effect (with three levels: 2016-2018) in all models. To facilitate convergence, chick age was standardized in all models.

After departure on autumn migration, for most chicks and some parents, only GSM-data were available for which the GPS-intervals varied between 1-6 hours. Moreover, because of the high speed during migratory flights, even if the birds were in the same flock, a small time-difference between the GPS-timestamp of the chick and the parent would already result in more than 10 meters distance between the birds. Therefore, we assessed whether chicks and parents may have been in contact during or after autumn migration by graphically exploring the cases where the distance between the nearest GPS locations was less than 25 km.

All statistical analyses were performed in program R (v. 4.4.1, R Core Team 2024) using package lme4 (Bates et al. 2015) to fit generalized linear mixed models and package DHARMa (Hartig 2022) to confirm homogeneity of residuals and lack of overdispersion. Model support was assessed on the basis of the Akaike information criterion adjusted for small sample sizes (Burnham and Anderson 2002). We selected the most parsimonious model as the model within 2 ΔAIC_c_ of the best-supported model that had the least parameters. Where applicable, post-hoc pairwise comparisons were performed using R-package emmeans (Lenth 2022).

## Results

### Visual feeding observations

The number of observed feedings of colour-ringed chicks after fledging strongly decreased with chick age (N=127, z=-8.87, p<0.001, Fig. 1a), with the oldest chick observed to be fed being 125 days old. While accounting for this age effect, the number of observed feedings did not significantly differ between female (N=40) and male chicks (N=46; z=0.56, p=0.58) nor between female (N=23) and male parents (N=34; z=1.28, p=0.20; Fig. S1). Of the 127 observed feedings, 70% occurred within 10 km from the natal colony, and the remaining 30% between 10 and 40 km distance from the colony (Fig. 1b, c). The distance from the colony at which chicks were fed increased with chick age (R^2^=0.33, F_1,125_=62.06, p<0.001, Fig. 1b).

**Fig. 1.**
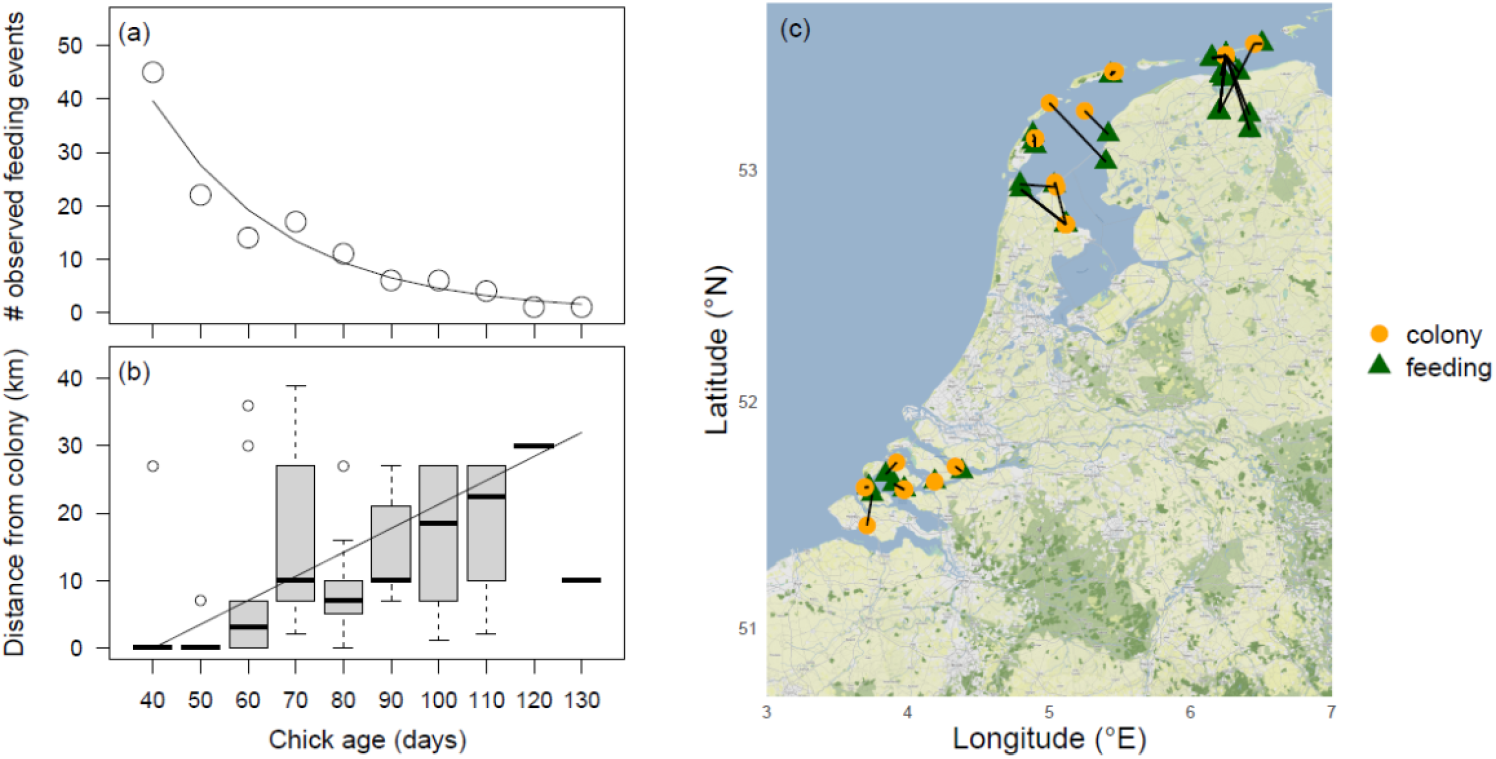
(a) The number of observations of colour-ringed chicks that were fed by an adult and (b) the distance from the colony at which these feedings took place in relation to chick age, along with a (c) map of the actual feeding locations connected by black lines to the colony locations where these chicks were born. Plotted data and model estimates are based on one randomly selected observation per known-age chick (N=127). To improve the readability of Fig. 1b, ages were binned, whereas in the analysis, chick age was treated as unbinned continuous variable. Boxes indicate the interquartile range (IQR), with the central line depicting the median, the whiskers extending to 1.5^*^IQR and open circles representing outliers

### Classification of begging behaviour

Begging behaviour was fairly accurately classified, with the highest classification performance achieved when using the longest segments of 1.6 s (F=0.89; Table S2). The other pooled behaviours were classified similarly well at 0.8 and 1.6 s (Table S2). As for this paper, the primary interest is in distinguishing begging and foraging, we trained and applied the random-forest model on 1.6 s acceleration data segments.

### GPS-based contact moments

The overall probability of contact (i.e. a distance of less than 10 m) between a GPS-tagged chick and its GPS-tagged parent was best explained by the model that included an effect of chick age and parent sex (Table S3). The probability of contact generally decreased with increasing chick age and mothers had somewhat more contact with their chick than fathers (Fig. 2, Table S4). Similar results were found for the probability of contact while the chick was classified to be begging or foraging (Table S3), although the slope of the age effect was opposite for foraging contact: older chicks were more likely to be foraging together with a parent than younger chicks (Tables S5, S6). However, the parameter estimates of the fixed effects were usually smaller than, or at most similar to, the estimated standard deviations (σ) for the random intercepts and slopes (Tables S4-S6). This indicates that the variation among chicks in the probability to have contact with their parent(s), and how this changed with age, was larger than what could be explained by the fixed effects (Figs. S2-S4). Indeed, the age at which a chick last had contact with its parent strongly varied, between 44 and 136 d (median: 88 d).

**Fig. 2.**
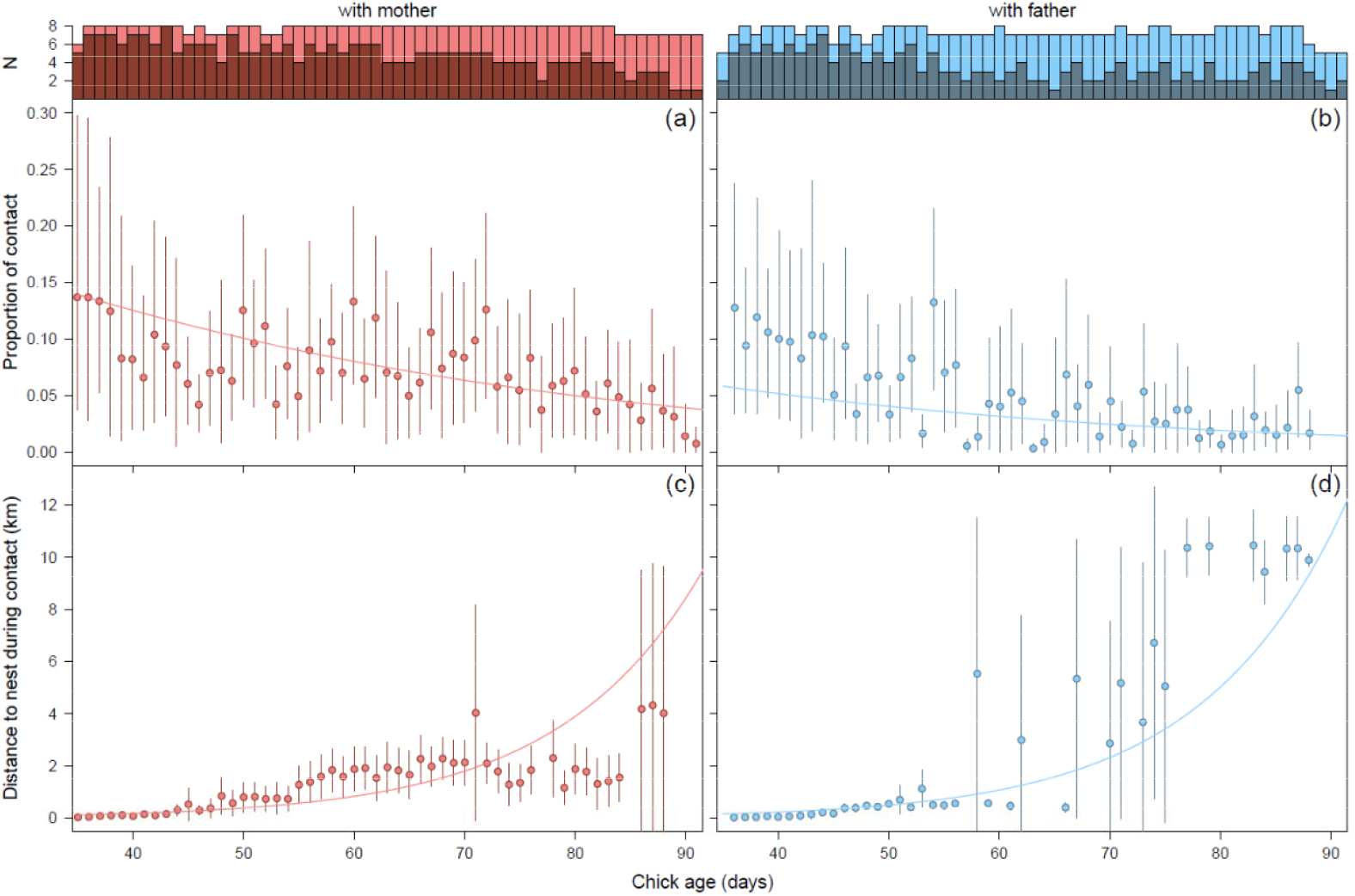
The proportion of time that a chick was in contact with its mother (a) or father (b) and the distance to the nest during contact with the mother (c) or father (d). Error bars reflect 95% CI, using the average proportion of time and distance to the nest during contact per day per chick as raw data. For the proportion data, bootstrapping was used to determine 95% CI’s. Sample sizes (number of chick-parent pairs) for the calculation of proportion of contact (total bars) and the distance to the nest during contact (darker colors) are shown above the panels. Only reasonably reliable estimates are plotted, i.e., estimates that were based on at least 6 different chick-parent pairs for the proportion of contact and at least 3 different chick-parent pairs for the distance to the nest during contact

Variation in distance to the nest during contact was also most parsimoniously explained by chick age and sex of the parent (Table S3). The distance to the nest during contact increased with chick age and contact with the father on average occurred further away from the nest than contact with the mother (Fig. 2; Table S7). When using a contact distance of 50 m instead of 10 m, the overall percentage of contact time (averaged over all chicks, irrespective of chick or parent sex) increased from 8.0% to 13.2% for 40 d old chicks and from 1.9% to 4.2% for 90 d old chicks. Model selection results were similar (Table S3 vs S8).

### Departure on autumn migration

Thirteen of the 15 GPS-tagged chicks departed on autumn migration, at ages varying between 108 and 139 d (median: 116 d). Except for one chick-parent pair, the last contact occurred prior to migration departure of either chick or parent: contact stopped 1 – 72 d (median: 28 d) prior to the chick’s departure and 0 – 70 d (median: 8 d) prior to the parent’s departure.

Of the 11 GPS-tagged chick-parent pairs where both chick and parent(s) departed on migration, only one chick departed on the same day as its parent (Fig. 3). Visual inspection of the timing and routes during their departing flights revealed that this chick (6295; a female) and its parent (6288; a male) were in the same flock (Fig. 4). However, while her father landed in the Kwade Hoek in the Dutch Delta, 6295 continued flying another 75 km to the Baai van Heist in northern Belgium. A few days later, they briefly met again at a roost along the Slikken van Flakkee, after which 6295 headed further south on 30 September, while her father stayed for another few days and departed on 2 October. So, even while they departed together from the north of The Netherlands, their connection was already lost in the south of The Netherlands.

**Fig. 3.**
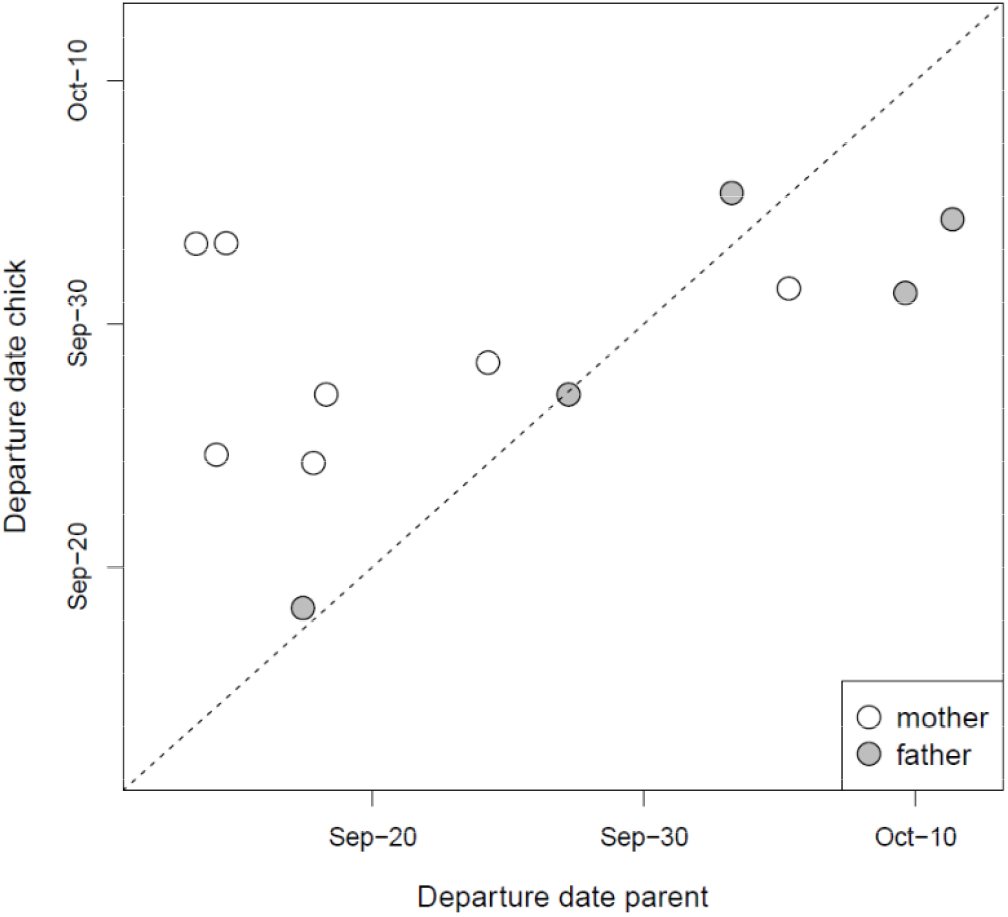
Autumn migration departure dates of spoonbill chicks and their parent(s) from The Netherlands. The dashed line (y = x) is added to visualize whether parents departed earlier (points above the line) or later (points below line) than their chicks

**Fig. 4.**
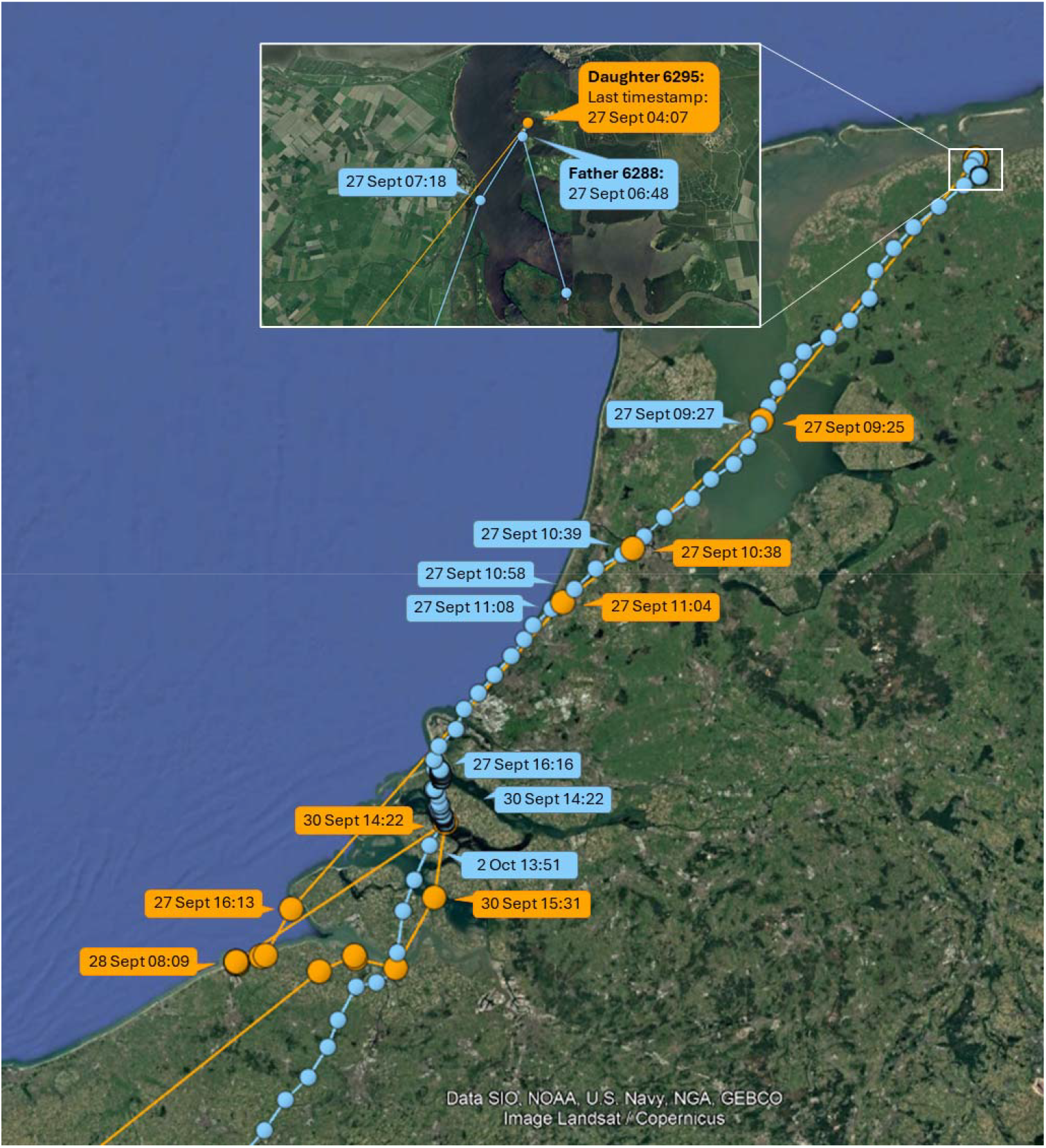
Detailed information about the timing and routes of the female chick 6295 (in orange) and her father 6288 (in blue) during the start of autumn migration from The Netherlands in 2016

The distance between the nearest (in time) GPS locations of the other 10 chick-parent pairs after their departure on migration, and of 6295-6288 after 30 September, was always >40 km, implying that chicks and parents did not meet again at later stages of the migration nor at the wintering sites.

## Discussion

Despite its potential role in affecting survival, habitat use and migration strategies of juvenile birds, post-fledging parental care is an understudied aspect of avian life histories due to the difficulty of following families over large distances. In this study, we investigated post-fledging parental care in Eurasian spoonbills in The Netherlands through a combination of visual feeding observations and GPS-tracking along with accelerometer-based classification of behaviours of parent(s) and their chick. We found that the number of feedings and the amount of contact that chicks still have with their parents strongly decreased with chick age and (usually) ends before the onset of autumn migration. The amount of post-fledging parental care received (and provided) strongly varied between chicks (and parents), with some evidence that mothers provided more care than fathers.

In our study, all visually observed feedings of juvenile spoonbills occurred within 40 km from the breeding colonies. This is similar to what was found in a Hungarian population, where the majority (86%) of fledged colour-ringed chicks were fed within 30 km of the breeding colonies (Pigniczki 2022). However, in this population, there were also two (out of 14) chicks observed being fed at respectively 111 km north and 140 km south of their colony. As observations along the Central-European flyway are scarce, this begged the question whether post-fledging parental care might sometimes extend into the migratory period. Yet, despite considerable observation efforts along the East-Atlantic flyway, with >10,000 observations of c. 4,000 juvenile birds from The Netherlands during their first autumn migration (i.e., observed south of The Netherlands during the months July-December) between 1994 and 2024, there was not a single observation of a juvenile being fed by a parent (Fig. 1). Also, for the GPS-tagged chicks and their parent(s), all contact moments were within 18 km from the colony, except for the contact between chick 6295 and father 6288 during their joint outbound migratory flight (Fig. 4). That some juveniles in Hungary were fed at considerably larger distances from the colony might be explained by potentially more variable food availability in this population, where spoonbills rely entirely on freshwater habitats that might dry out or become densely vegetated in the course of the summer. Contrastingly, in our dataset, the majority of observed feedings came from chicks from the Wadden Sea area or the Dutch Delta, where many marine and brackish foraging grounds remain available throughout the breeding season.

While the GPS-data revealed that female parents had more contact with their chick after fledging than male parents, female parents were not more often observed feeding a chick (N=23) than male parents (N=34). In fact, although the difference was not statistically significant, the visual feeding observations even pointed in the opposite direction. In absence of a strong bias in the percentage of females among the visually or molecularly sexed adults in the entire resighting dataset (48%), there remain several explanations for this apparently contrasting result. Firstly, contact does not necessarily mean that a chick is fed. However, the GPS-tagged chicks were also more likely to be begging at their mother than at their father, which may be assumed to reflect the probability of being fed. Alternatively, the estimated parental sex effect in the GPS-data may not be very representative as it strongly driven by the data of the two chicks of which both parents were GPS-tagged. In both cases, the mother had more contact with the chick, overall and while the chick was begging or foraging, than the father (Figs. S2-S4). But with a sample size of just two, this may not necessarily be a general pattern. Among chicks, there was considerable variation in the amount of contact they still had with their parent, whether it was with their father or their mother (Figs. S2-S4). Consequently, if we would treat the two chicks of which both parents were GPS-tagged as four unique chick-parent pairs (i.e., by using chick-parent pair instead of chick as random effect), the effect of sex of the parent was no longer supported (Table S9).

We used contact between GPS-tagged chicks and parent(s) as an indication of post-fledging parental care. However, it is uncertain, at least when it comes to contact while the chick was resting or foraging (the chick was on average only begging during c. 3% of its contact time), whether this contact was ‘deliberate’ (i.e. that the parent was actively protecting or guiding the chick, or the chick actively sought out or followed its parent) or ‘coincidental’. The latter could be caused by the sociality of spoonbills, irrespective of their relatedness. Shortly after fledging, social structuring occurs between roosts near the colony and adjacent foraging grounds in the Wadden Sea. Later on, spoonbills from multiple colonies congregate in larger groups, where they moult and prepare for migration (e.g., fuel and seek companions to depart with; de Boer et al. 2024; Henriques et al. 2025). That at least part of the contact, particularly at older ages, may be coincidental is suggested by the fact that six chick-parent pairs showed a long period without contact after which there was contact again (Figs. 3, S2). In four of these cases (6295-6288, 6298.1-6118, 6304-763 and 6358.1-6066), this was caused by the fact that the parent (in all four cases the father) no longer returned to the colony and at a certain point, the chick dispersed away from the colony and met the parent again (Fig. 5). In the other two cases (6315-6289 and 6385-6291), the chick was the first to move away from the colony, whereas the parent (in both cases the mother) followed later (Fig. 5).

**Fig. 5.**
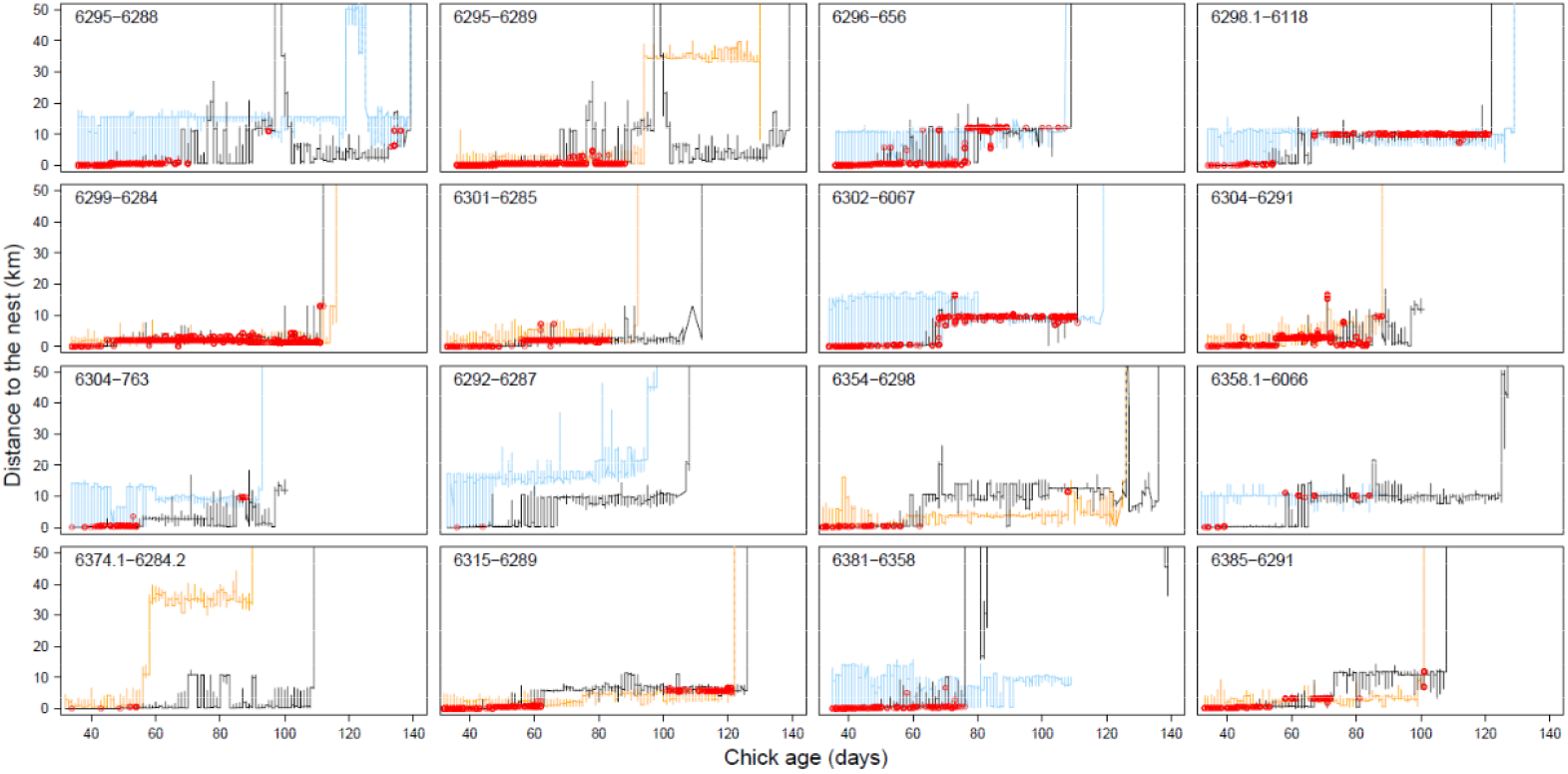
Distance to the nest of chick (in black) and parent (fathers in blue, mothers in orange), with the open red circles indicating the contact moments. Here the raw data are plotted, i.e. also the days with less than 130 registered GPS locations for both chick and parent which was the criterion to be included in the analyses. Some distances to the nest are not plotted as they exceeded 50 km, including the last vertical lines representing the outbound migratory flights. All contact moments are plotted except for the single contact moment between 6295 and 6288 in the Dutch Delta. Although 6295 and 6288 were also in the same flock during their outbound migratory flight, this was not estimated as contact as the distance between their locations was larger than 10 m due to the time difference (varying between 1 and 4 minutes) between the timestamps of chick and parent combined with the high speed during migration (see Fig. S4). Furthermore, although it may appear that 6354 attempted to depart with his mother 6298, this was not the case: the departure location, time and routes of their flights were different

Our results indicate that the amount of post-fledging parental care received (and provided) is highly variable among chicks (and parents). First of all, it should be realized that in most cases, we only have information on the contact between the chick and one of its parents. Therefore, it might be possible that the untagged parent compensates for the lack of care provided by some of the GPS-tagged parents. Yet, in the two cases where both parents were GPS-tagged, both provided post-fledging parental care, making this a rather unlikely scenario. The variable care provided was also not explained by the fact that some parents had to divide their care over multiple chicks: among the four GPS-tagged chicks that had a younger or older sibling (6298.1, 6299, 6292 and 6381), three of them received hardly any post-fledging parental care, whereas 6299 was among the chicks that received the most care. Similarly, among the ten GPS-tagged chicks that were the only chick in the nest that survived until fledging, the amount of care received also varied considerably.

It is also possible that parents compensate for differences in the amount of care they already provided prior to fledging. During incubation and nest-bound chick-rearing, females and males equally shared nest attendance duties, but females spent more time foraging (Lok et al. 2024). As a result, female parents may be expected to contribute less to post-fledging parental care than males, but this was not the case. In addition to potential sex-specific energetic considerations, parents are known to vary in terms of migration distance (Lok et al. 2017), with long-distance migrants potentially being more constrained in terms of the amount of care they can afford to provide while they also invest in moult and pre-migratory fuelling.

Rather than (or in addition to) being caused by variation in the availability or energetic state of the parents, variation in post-fledging parental care could be caused by variation in the energetic requirements of the chicks. When food availability is low, chicks may fledge in poorer condition and have more difficulties in finding food for themselves, therefore begging their parents to feed them for a longer period of time in order to survive. On the other hand, under poor food conditions, parents may decide to secure their own survival and future reproductive prospects at the cost of the survival of their current young, as is expected and commonly observed in long-lived species (Stearns 1992). That the latter might play a role in explaining the observed variation is supported by the fact that in 2017, the year in which hardly any post-fledging parental care was provided, breeding success (unpublished data) as well as estimated prey ingestion rates during late summer in the Wadden Sea (Lok et al. 2023) were relatively low. Although estimated prey ingestion rates in the Wadden Sea were similarly low in 2018, this year’s summer was extremely dry, causing the freshwater lake Westerplas on Schiermonnikoog to dry out nearly completely with the fish being trapped in a very small layer of water: a foraging feast for spoonbills (PdG, pers. obs.).

Rather than by energetic aspects, the observed inter-family variation in post-fledging care could also be caused by different processes. It is now well established that birds often exhibit consistent between-individual differences in boldness and explorativeness across different life stages and environmental contexts (Réale et al. 2007). Some chicks may simply be more demanding than others, persistently following their parents for weeks. Similarly, some parents may be more caring than others, regardless of their energetic state.

Despite the considerable variation in the extent of contact prior to departure, in 10 of the 11 GPS-tagged families where both chick and parent departed on autumn migration, the chick departed on a different day than (i.e. independently from) its parent. In the first half of the migratory season (until 2 October), juveniles usually departed later than their GPS-tagged parent, whereas later on, the juveniles (N=3) departed earlier. There was only one case where the chick departed in the same flock as its parent, but they did not stop at the same place, hence contact was already lost prior to leaving The Netherlands (Fig. 3). Moreover, prior to the two days preceding their joint southward departure, the chick and its parent had not been in contact for 36 days (Figs. 3, S2). This suggests that it was not the bond between chick and parent, but rather coincidence due to spoonbills socially migrating in (large) flocks, that caused them to be in the same flock during their departing flight. Therefore, our results suggest that, unlike geese, swans, cranes and several other species (Scott 1980; Black and Owen 1989; Earnst and Bart 1991; Alonso et al. 2004; Palacín et al. 2011; Chetverikova et al. 2017; Byholm et al. 2022), spoonbill chicks and parents do not deliberately migrate together. Apparently, the costs of prolonged care by the parents during migration are higher than the benefits gained from it by either the juveniles (e.g., by receiving food and/or being guided during migration) or the parents (e.g., through increased dominance status). However, as spoonbills usually migrate in large flocks, consisting of both adult and juvenile birds (Piersma et al. 2022), juveniles may use the information from other experienced adults to learn suitable migration routes, to find staging and wintering sites, and to exploit local resources across a variety of locations and environments – often very different from the environment in which they were born and cared for.

To conclude, both parents contribute to post-fledging parental care, with considerable variation in the amount of care provided. It remains unclear whether this variation is caused by differences in the demands of chicks to still receive post-fledging parental care, the demands of the parents, or by different processes entirely. The amount of parental care strongly decreases with chick age and (usually) ends several weeks before departing on autumn migration with other conspecifics.

## Acknowledgments

We thank Natuurmonumenten for permission to work in national park Schiermonnikoog, Yvonne Verkuil for molecular sexing, Willem Bouten for technical assistance on UvA-BiTS, Iris Kromhout Van der Meer, Marnix Bosma, Remco Rood and many other volunteers for their assistance in the field and Werkgroep Lepelaar and voluntary ring-readers for ringing and resighting (and observing the behaviour and sex of) colour-ringed spoonbills.

## Author contributions

TL (Conceptualization, Data curation, Formal analysis, Funding acquisition, Investigation, Methodology, Project administration, Visualization, Writing—original draft, Writing—review and editing), PdG (Investigation, Methodology, Project administration, Writing—review and editing), ER (Data curation, Methodology, Resources, Writing—review and editing), TP (Conceptualization, Funding acquisition, Methodology, Project administration, Supervision, Writing—review and editing) and WV (Conceptualization, Supervision, Visualization, Writing—review and editing).

## Data availability

The data and R code to reproduce the analyses and results reported in this article are available online (Lok 2025) and maintained at GitHub (https://github.com/tamarlok/postfledging_care).

## Declarations

### Ethical approval

This study was conducted under license numbers D6548 and AVD105002016446 following the Dutch Animal Welfare Act and the ethical standards of the Dutch Centre for Avian Migration & Demography.

---

**Photo 1.**
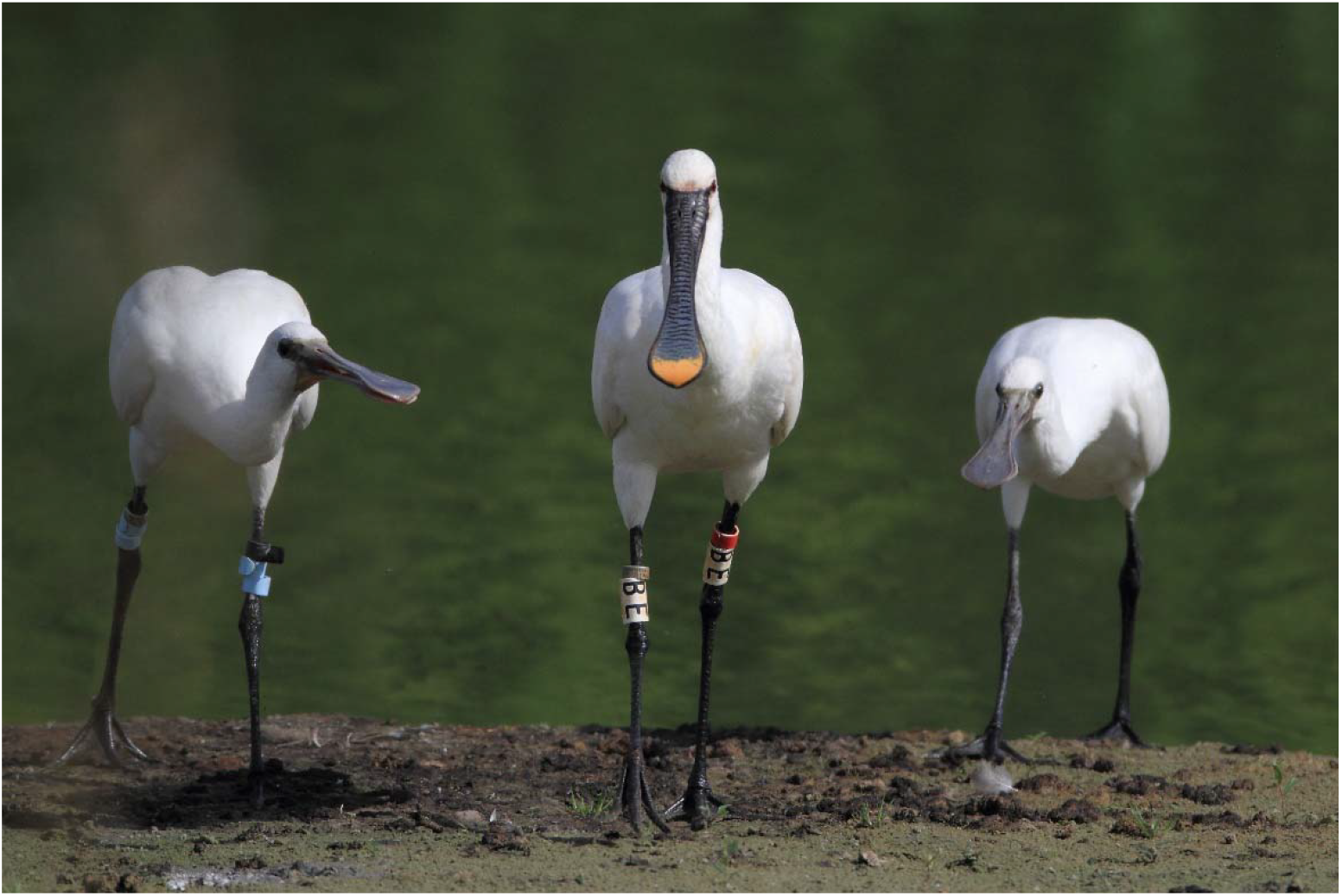
Adult spoonbill followed by two begging chicks during the post-fledging period at Braakman, Hoek (51°196□N 3°445□E), 16 August 2015. The chick was colour-ringed on 6 July 2015 in the colony at Van Cittershaven, Sloegebied (51°263□N 3°433□E), 13 km from the location of this photo. Photographer: Gab de Croock

## References

Alonso JC, Bautista LM, Alonso JA (2004) Family-based territoriality vs flocking in wintering common cranes Grus grus. J Avian Biol 35:434–444. 10.1111/j.0908-8857.2004.03290.x

Ashmole NP, Tovar SH (1968) Prolonged parental care in royal terns and other birds. Auk 85:90–100. 10.2307/4083627

Bates D, Maechler M, Bolker B, Walker S (2015) Fitting linear mixed-effects models using lme4. J Stat Softw 67:1–48. 10.18637/jss.v067.i01

Black JM, Owen M (1989) Parent-offspring relationships in wintering barnacle geese. Anim Behav 37:187–198. 10.1016/0003-3472(89)90109-7

Bouten W, Baaij EW, Shamoun-Baranes J, Camphuysen KCJ (2013) A flexible GPS tracking system for studying bird behaviour at multiple scales. J Ornithol 154:571–580. 10.1007/s10336-012-0908-1

Burnham K, Anderson D (2002) Model selection and multi-model inference: a practical information-theoretic approach, 2nd edn. Springer-Verlag, New York

Byholm P, Beal M, Isaksson N, Lötberg U, Åkesson S (2022) Paternal transmission of migration knowledge in a long-distance bird migrant. Nat Commun 13:1566. 10.1038/s41467-022-29300-w

Chetverikova R, Babushkina O, Galkina S, Shokhrin V, Bojarinova J (2017) Special case among passerine birds: long-tailed tits keep family bonds during migration. Behav Ecol Sociobiol 71:40. 10.1007/s00265-017-2268-6

Cramp S (1994) The birds of the Western Palearctic. Oxford University Press, Oxford

de Boer AP, Vansteelant WMG, Piersma T (2024) Primary moult of Eurasian spoonbills Platalea l. leucorodia in the Wadden Sea in relation to age, breeding and migration. Ardea 112:217–227. 10.5253/arde.2023.a19

Earnst SL, Bart J (1991) Costs and benefits of extended parental care in Tundra Swans Cygnus columbianus columbianus. Wildfowl Special Issue 1:260–267

Ersoy S, Maag N, Boehly T, Boucherie PH, Bugnyar T (2021) Sex-specific parental care during postfledging in common ravens. Anim Behav 181:95–103. 10.1016/j.anbehav.2021.09.004

Fijn RC, Kemper J (2023) Sandwich tern feeds juvenile on wintering grounds in southern Namibia. Ardea 111:549–553. 10.5253/arde.2023.a2

Fridolfsson AK, Ellegren H (1999) A simple and universal method for molecular sexing of non-ratite birds. J Avian Biol 30:116–121. 10.2307/3677252

Grüebler MU, Naef-Daenzer B (2010) Survival benefits of post-fledging care: experimental approach to a critical part of avian reproductive strategies. J Anim Ecol 79:334–341. 10.1111/j.1365-2656.2009.01650.x

Henriques M, Piersma T, Vansteelant WMG, de Goeij P, Lok T (2025) A time and a place for everything: Eurasian Spoonbills divide spring and summer activities across different areas in the eastern Dutch Wadden Sea. Ardea 113:3–20. 10.5253/arde.2024.a24

Koenig WD, Dickinson JL (2004) Ecology and evolution of cooperative breeding in birds. Cambridge University Press, Cambridge

Krijgsveld KL, Dijkstra C, Visser GH, Daan S (1998) Energy requirements for growth in relation to sexual size dimorphism in marsh harrier Circus aeruginosus nestlings. Physiol Zool 71:693–702. 10.1086/515983

Lagarde F, Piersma T (2021) Vocal signalling by Eurasian Spoonbills Platalea leucorodia in flocks before migratory departure. Ardea 109:243–250. 10.5253/arde.v109i3.a8

Lenth RV (2022) emmeans: Estimated Marginal Means, aka Least-Squares Means. R package version 1.7.2, https://cran.r-project.org/web/packages/emmeans

Lok T (2025) R code and dataset for: Eurasian Spoonbill chicks receive parental care up to several months after fledging, but not into migration. Behav Ecol Sociobiol. [Computer software]. Zenodo. 10.5281/zenodo.16961361

Lok T, Overdijk O, Piersma T (2014) Interpreting variation in growth of Eurasian Spoonbill chicks: disentangling the effects of age, sex and environment. Ardea 102:181–194. 10.5253/arde.v102i2.a8

Lok T, Overdijk O, Tinbergen JM, Piersma T (2013) Seasonal variation in density dependence in age-specific survival of a long-distance migrant. Ecology 94:2358–2369. 10.1890/12-1914.1

Lok T, van der Geest M, Bom RA, de Goeij P, Piersma T, Bouten W (2023) Prey ingestion rates revealed by back-mounted accelerometers in Eurasian spoonbills. Anim Biotelem 11:5. 10.1186/s40317-022-00315-w

Lok T, van der Geest M, de Goeij P, Rakhimberdiev E, Piersma T (2024) Sex-specific nest attendance rhythm and foraging habitat use in a colony-breeding waterbird. Behav Ecol 35:arae020. 10.1093/beheco/arae020

Lok T, Veldhoen L, Overdijk O, Tinbergen JM, Piersma T (2017) An age-dependent fitness cost of migration? Old trans-Saharan migrating spoonbills breed later than those staying in Europe, and late breeders have lower recruitment. J Anim Ecol 86:998–1009. 10.1111/1365-2656.12706

López-Idiáquez D, Vergara P, Fargallo JA, Martínez-Padilla J (2018) Providing longer post-fledging periods increases offspring survival at the expense of future fecundity. PLoS ONE 13:e0203152. 10.1371/journal.pone.0203152

Mellink E, Castillo-Guerrero JA, Peñaloza-Padilla E (2014) Development of diving abilities by fledgling Brown Boobies (Sula leucogaster) in the Central Gulf of California, México. Waterbirds 37:451–456. 10.1675/063.037.0414

Meyburg B-U, Meyburg C, Mizera T, Maciorowski G, Kowalski J (2005) Family break up, departure, and autumn migration in Europe of a family of greater spotted eagles (Aquila clanga) as reported by satellite telemetry. J Raptor Res 39:462–466

Naef-Daenzer B, Grüebler MU (2016) Post-fledging survival of altricial birds: ecological determinants and adaptation. J Field Ornithol 87:227–250. 10.1111/jofo.12157

Palacín C, Alonso JC, Alonso JA, Magaña M, Martín CA (2011) Cultural transmission and flexibility of partial migration patterns in a long-lived bird, the great bustard Otis tarda. J Avian Biol 42:301–308. 10.1111/j.1600-048x.2011.05395.x

Phillips RA, Xavier JC, Croxall JP (2003) Effects of satellite transmitters on albatrosses and petrels. Auk 120:1082. 10.1093/auk/120.4.1082

Piersma T, de Goeij P, Bouten W, Zuhorn C (2022) Sinagote. The biography of a spoonbill. Lynx Edicions, Barcelona

Pigniczki C (2022) Observations on parental care of the Eurasian Spoonbill during the post-fledging dispersal. Ornis Hung 30:146–157. 10.2478/orhu-2022-0011

R Core Team (2024) R: A language and environment for statistical computing. In. R Foundation for Statistical Computing, Vienna, Austria, http://www.R-project.org

Réale D, Reader SM, Sol D, McDougall PT, Dingemanse NJ (2007) Integrating animal temperament within ecology and evolution. Biol Rev 82:291–318. 10.1111/j.1469-185X.2007.00010.x

Rehling A, Spiller I, Krause ET, Nager RG, Monaghan P, Trillmich F (2012) Flexibility in the duration of parental care: zebra finch parents respond to offspring needs. Anim Behav 83:35–39. 10.1016/j.anbehav.2011.10.003

Riou S, Chastel O, Hamer KC (2012) Parent–offspring conflict during the transition to independence in a pelagic seabird. Behav Ecol 23:1102–1107. 10.1093/beheco/ars079

Scott DK (1980) Functional aspects of prolonged parental care in Bewicks swans. Anim Behav 28:938–952. 10.1016/s0003-3472(80)80156-4

Stearns SC (1992) The evolution of life histories. Oxford University Press, Oxford

Whiten A (2019) Cultural evolution in animals. Annu Rev Ecol Evol S 50:27–48

