## Supporting Information for "Eurasian Spoonbill chicks receive parental care up to several months after fledging, but not into migration"

accompanying the paper

Behavioral Ecology and Sociobiology

Tamar Lok[1,2,3*], Petra de Goeij[4], Eldar Rakhimberdiev[5], Theunis Piersma[1,3,4] & Wouter Vansteelant[1,4,5]

[1] Rudi Drent Chair in Global Flyway Ecology, Conservation Ecology Group, Groningen Institute for Evolutionary Life Sciences, University of Groningen, Nijenborgh 9, 9747 AG Groningen, Netherlands
[2] EcoMoves, Wilhelminalaan 78, 1791 AP Den Burg, The Netherlands
[3] Department of Coastal Systems, NIOZ Royal Netherlands Institute for Sea Research, P.O. Box 59, 1790 AB, Den Burg, Texel, The Netherlands
[4] BirdEyes, Centre for Global Ecological Change at the Faculties of Science & Engineering and Campus Fryslân, University of Groningen, Wirdumerdijk 34, 8911 CE, Leeuwarden, The Netherlands
[5] Department of Theoretical and Computational Ecology, Institute for Biodiversity and Ecosystems Dynamics, University of Amsterdam, Science Park 904, 1098 XH Amsterdam, the Netherlands

**Table S1.**

Information about GPS-tagged parents and chicks, nests and joined GPS data. The number of days with nearly complete data used in the analyses is specified separately for modelling overall contact and (between brackets) behaviour-specific contact.

| Parent info | | | Chick info | | | | | | Nest info | | | GPS data info | |
|---|---|---|---|---|---|---|---|---|---|---|---|---|---|
| ID | Sex | Deploy-ment date | ID | Sex | Deployment date | Deploy-ment age (d) | Chick order | Mortality date | Latitude | Longitude | # fledg-lings | # days used in the analyses | Last date with (some) joined data |
| 6288 | m | 12/05/2016 | 6295 | f | 15/06/2016 | 35 | 1 | | 53.4865 | 6.23798 | 1 | 97 (89) | 31/12/2016 |
| 6289 | f | 15/05/2016 | 6295 | f | 15/06/2016 | 35 | 1 | | 53.4865 | 6.23798 | 1 | 98 (91) | 31/12/2016 |
| 656 | m | 27/04/2016 | 6296 | m | 06/07/2016 | 35 | 1 | 03/10/2016 | 53.49024 | 6.26037 | 1 | 51 (48) | 03/10/2016 |
| 6118 | m | 30/04/2014 | 6298.1 | m | 07/07/2016 | 33 | 1 | 27/10/2016 | 53.48836 | 6.25927 | 2 | 86 (85) | 28/10/2016 |
| 6284 | f | 28/04/2016 | 6299 | f | 14/07/2016 | 33 | 1 | | 53.49356 | 6.27743 | 2 | 77 (75) | 31/12/2016 |
| 6285 | f | 02/05/2016 | 6301 | f | 15/07/2016 | 32 | 1 | 01/11/2016 | 53.49354 | 6.27751 | 1 | 63 (61) | 01/11/2016 |
| 6067 | m | 13/05/2014 | 6302 | f | 15/07/2016 | 33 | 1 | 25/11/2017 | 53.4864 | 6.26722 | 1 | 75 (70) | 29/12/2016 |
| 6291 | f | 16/05/2016 | 6304 | m | 28/07/2016 | 33 | 1 | 05/10/2016 | 53.48679 | 6.26757 | 1 | 65 (65) | 04/10/2016 |
| 763 | m | 25/05/2012 | 6304 | m | 28/07/2016 | 33 | 1 | 05/10/2016 | 53.48679 | 6.26757 | 1 | 65 (65) | 04/10/2016 |
| 6298 | f | 06/05/2017 | 6354 | f | 11/06/2017 | 31 | 1 | | 53.4862 | 6.25041 | 1 | 85 (81) | 31/12/2017 |
| 6066 | m | 05/05/2014 | 6358.1 | f | 12/06/2017 | 31 | 1 | 27/02/2018 | 53.48378 | 6.24165 | 1 | 54 (53) | 31/12/2017 |
| 6287 | m | 03/05/2016 | 6292 | f | 13/07/2017 | 32 | 2 | 06/10/2017 | 53.47796 | 6.24427 | 2 | 67 (57) | 06/10/2017 |
| 6284.2 | f | 01/05/2017 | 6374.1 | m | 17/07/2017 | 31 | 1 | 03/01/2018 | 53.48367 | 6.23753 | 1 | 74 (65) | 31/12/2017 |
| 6288 | m | 12/05/2016 | 6374.1 | m | 17/07/2017 | 31 | 1 | 03/01/2018 | 53.48367 | 6.23753 | 1 | 0 (0) | 31/12/2017 |
| 6358 | m | 14/05/2018 | 6381 | m | 21/06/2018 | 34 | 2 | | 53.48696 | 6.25897 | 2 | 27 (23) | 31/12/2018 |
| 6284.2 | f | 01/05/2017 | 1607 | m | 25/06/2018 | 30 | 1 | | 53.47805 | 6.24248 | 1 | 0 (0) | 02/07/2018 |
| 6289 | f | 15/05/2016 | 6315 | m | 25/06/2018 | 31 | 1 | | 53.48611 | 6.2353 | 1 | 90 (79) | 31/12/2018 |
| 6288 | m | 12/05/2016 | 6315 | m | 25/06/2018 | 31 | 1 | | 53.48611 | 6.2353 | 1 | 0 (0) | - |
| 6291 | f | 16/05/2016 | 6385 | m | 10/07/2018 | 32 | 1 | | 53.48726 | 6.26056 | 1 | 70 (50) | 31/12/2018 |

**Table S2.**

F-measures for pooled behaviours, where ‘forage’ includes the behaviours ‘search’, ‘handle’ and ‘ingest’ from Lok et al. 2023, ‘rest’ includes ‘sit’ and ‘stand’, also when the bird is moving its body while standing (i.e. when shaking feathers, preening or drinking) and ‘fly’ includes active and passive flight. The random-forest model was trained on the accelerometer dataset (20 Hz 3D data) with unpooled annotated behaviours ‘search’, ‘handle’, ‘ingest’, ‘sit’, ‘stand’, ‘fly-active’, ‘fly-passive’, ‘walk’ and ‘beg’. Reported values are means calculated from 100 runs of 70% training and 30% test data, randomly sampled from the annotated dataset.

| | **Segment length (s)** | | |
|---|---|---|---|
| | **0.4** | **0.8** | **1.6** |
| **forage** | 0.95 | 0.96 | 0.96 |
| **walk** | 0.53 | 0.60 | 0.61 |
| **fly** | 0.97 | 0.97 | 0.98 |
| **rest** | 0.97 | 0.96 | 0.95 |
| **beg** | 0.79 | 0.86 | 0.89 |

**Table S3.**

Model selection results. The most parsimonious models are indicated in bold. "age" refers to age of the chick and "c" to the intercept-only "constant" model. For modelling contact probabilities, year was always included as a categorical fixed effect.

| Model | K | Δ-2logL | $\Delta AIC_c$ | Akaike weight |
|---|---|---|---|---|
| *(a) Total contact* | | | | |
| **sex$_{parent}$ + age** | **8** | **0.79** | **0.00** | **0.64** |
| sex$_{chick}$ + sex$_{parent}$ + age | 9 | 0.00 | 1.21 | 0.35 |
| sex$_{parent}$ | 7 | 11.46 | 8.67 | 0.01 |
| sex$_{chick}$ + sex$_{parent}$ | 8 | 10.67 | 9.88 | 0.00 |
| age | 7 | 601.05 | 598.26 | 0.00 |
| sex$_{chick}$ + age | 8 | 599.56 | 598.76 | 0.00 |
| c | 6 | 611.58 | 606.79 | 0.00 |
| sex$_{chick}$ | 7 | 610.04 | 607.25 | 0.00 |
| *(b) Contact while begging* | | | | |
| **sex$_{parent}$ + age** | **8** | **0.71** | **0.00** | **0.65** |
| sex$_{chick}$ + sex$_{parent}$ + age | 9 | 0.00 | 1.29 | 0.34 |
| sex$_{chick}$ + age | 8 | 10.53 | 9.81 | 0.00 |
| sex$_{parent}$ | 7 | 13.55 | 10.83 | 0.00 |
| age | 7 | 13.99 | 11.28 | 0.00 |
| sex$_{chick}$ + sex$_{parent}$ | 8 | 12.95 | 12.24 | 0.00 |
| sex$_{chick}$ | 7 | 23.55 | 20.84 | 0.00 |
| c | 6 | 26.77 | 22.06 | 0.00 |
| *(c) Contact while foraging* | | | | |
| **sex$_{parent}$ + age** | **8** | **0.57** | **0.00** | **0.60** |
| sex$_{chick}$ + sex$_{parent}$ + age | 9 | 0.00 | 1.43 | 0.29 |
| sex$_{parent}$ | 7 | 6.80 | 4.23 | 0.07 |
| sex$_{chick}$ + sex$_{parent}$ | 8 | 6.06 | 5.49 | 0.04 |
| age | 7 | 35.64 | 33.07 | 0.00 |
| sex$_{chick}$ + age | 8 | 34.77 | 34.20 | 0.00 |
| c | 6 | 42.77 | 38.20 | 0.00 |
| sex$_{chick}$ | 7 | 41.54 | 38.97 | 0.00 |
| *(d) Distance from nest during contact* | | | | |
| **sex$_{parent}$ + age** | **7** | **0.13** | **0.00** | **0.72** |
| sex$_{chick}$ + sex$_{parent}$ + age | 8 | 0.00 | 1.87 | 0.28 |
| sex$_{parent}$ | 6 | 25.71 | 23.57 | 0.00 |
| sex$_{chick}$ + sex$_{parent}$ | 7 | 25.54 | 25.41 | 0.00 |
| age | 6 | 38.90 | 36.76 | 0.00 |
| sex$_{chick}$ + age | 7 | 38.78 | 38.65 | 0.00 |
| c | 5 | 64.26 | 60.13 | 0.00 |
| sex$_{chick}$ | 6 | 64.13 | 62.00 | 0.00 |

**Table S4.**

Parameter estimates (on logit-scale) of the global model for overall contact probability.

| Parameter | Estimate | s.e. |
|---|---|---|
| *Fixed effects* | | |
| Intercept[1] | -2.06 | 0.36 |
| $Sex_{parent}$ (male) | -1.01 | 0.04 |
| $Sex_{chick}$ (male) | -0.46 | 0.43 |
| $Age_{chick}$ | -0.64 | 0.16 |
| Year (2017) | -3.44 | 0.57 |
| Year (2018) | -0.67 | 0.60 |
| *Random effects* | | |
| $\sigma_{Intercept(chickID)}$ | 0.81 | - |
| $\sigma_{Age(chickID)}$ | 0.59 | - |
| Cor($Intercept_{chickID}$, $Age_{chickID}$) | 0.28 | - |

[1]Reference levels are female for $Sex_{parent}$ and $Sex_{chick}$ and 2016 for Year.

**Table S5.**

Parameter estimates (on logit-scale) of the global model for contact probability while the chick is begging.

| Parameter | Estimate | s.e. |
|---|---|---|
| *Fixed effects* | | |
| Intercept[1] | -6.17 | 0.22 |
| $Sex_{parent}$ (male) | -0.53 | 0.17 |
| $Sex_{chick}$ (male) | -0.21 | 0.25 |
| $Age_{chick}$ | -0.98 | 0.24 |
| Year (2017) | -2.11 | 0.38 |
| Year (2018) | 0.16 | 0.31 |
| *Random effects* | | |
| $\sigma_{Intercept(chickID)}$ | 0.52 | - |
| $\sigma_{Age(chickID)}$ | 0.74 | - |
| Cor($Intercept_{chickID}$, $Age_{chickID}$) | 0.92 | - |

[1]Reference levels are female for $Sex_{parent}$ and $Sex_{chick}$ and 2016 for Year.

**Table S6.**

Parameter estimates (on logit-scale) of the global model for contact probability while the chick is foraging.

| Parameter | Estimate | s.e. |
|---|---|---|
| *Fixed effects* | | |
| Intercept[1] | -4.76 | 0.66 |
| $Sex_{parent}$ (male) | -1.49 | 0.28 |
| $Sex_{chick}$ (male) | -0.78 | 0.96 |
| $Age_{chick}$ | 0.65 | 0.22 |
| Year (2017) | -3.67 | 1.05 |
| Year (2018) | -1.00 | 1.18 |
| *Random effects* | | |
| $\sigma_{Intercept(chickID)}$ | 1.41 | - |
| $\sigma_{Age(chickID)}$ | 0.66 | - |
| Cor($Intercept_{chickID}$, $Age_{chickID}$) | 0.27 | - |

[1]Reference levels are female for $Sex_{parent}$ and $Sex_{chick}$ and 2016 for Year.

**Table S7.**

Parameter estimates of the global model for distance to the nest (in km, on log-scale) during contact.

| Parameter | Estimate | s.e. |
|---|---|---|
| *Fixed effects* | | |
| Intercept[1] | 0.97 | 0.29 |
| $Sex_{parent}$ (male) | 0.25 | 0.04 |
| $Sex_{chick}$ (male) | -0.13 | 0.37 |
| $Age_{chick}$ | 1.87 | 0.23 |
| *Random effects* | | |
| $\sigma_{Intercept(chickID)}$ | 0.81 | - |
| $\sigma_{Age(chickID)}$ | 0.83 | |
| $\sigma_{Residual}$ | 1.05 | - |
| Cor($Intercept_{chickID}$, $Age_{chickID}$) | 0.53 | - |

[1]Reference levels are female for $Sex_{parent}$ and $Sex_{chick}$ and 2016 for Year.

**Table S8.**

Model selection results for probability of contact, (a) overall, (b) while the chick is begging or (c) while the chick is foraging when using a contact distance of 50 m instead of 10 m (for which results are shown in Table 1). The most parsimonious models are indicated in bold. “age” refers to age of the chick and “c” to the intercept-only “constant” model. For modelling contact probabilities, year was always included as a categorical fixed effect.

| Model | K | Δ-2logL | $\Delta AIC_c$ | Akaike weight |
|---|---|---|---|---|
| *(a) Total contact* | | | | |
| **$sex_{parent}$ + age** | **8** | **0.98** | **0.00** | **0.48** |
| $sex_{chick}$ + $sex_{parent}$ + age | 9 | 0.00 | 1.02 | 0.29 |
| $sex_{parent}$ | 7 | 5.40 | 2.42 | 0.14 |
| $sex_{chick}$ + $sex_{parent}$ | 8 | 4.41 | 3.43 | 0.09 |
| age | 7 | 758.46 | 755.47 | 0.00 |
| $sex_{chick}$ + age | 8 | 757.33 | 756.34 | 0.00 |
| c | 6 | 762.89 | 757.91 | 0.00 |
| $sex_{chick}$ | 7 | 761.76 | 758.78 | 0.00 |
| *(b) Contact while begging* | | | | |
| $sex_{chick}$ + $sex_{parent}$ + age | 9 | 0.00 | 0.00 | 0.58 |
| **$sex_{parent}$ + age** | **8** | **2.79** | **0.79** | **0.39** |
| $sex_{chick}$ + $sex_{parent}$ | 8 | 9.30 | 7.30 | 0.02 |
| $sex_{parent}$ | 7 | 12.08 | 8.08 | 0.01 |
| $sex_{chick}$ + age | 8 | 27.92 | 25.92 | 0.00 |
| age | 7 | 32.62 | 28.62 | 0.00 |
| $sex_{chick}$ | 7 | 37.40 | 33.39 | 0.00 |
| c | 6 | 42.06 | 36.06 | 0.00 |
| *(c) Contact while foraging* | | | | |
| $sex_{chick}$ + $sex_{parent}$ + age | 9 | 0.00 | 0.00 | 0.40 |
| **$sex_{parent}$ + age** | **8** | **2.65** | **0.65** | **0.29** |
| $sex_{chick}$ + $sex_{parent}$ | 8 | 3.60 | 1.60 | 0.18 |
| $sex_{parent}$ | 7 | 6.09 | 2.09 | 0.14 |
| $sex_{chick}$ + age | 8 | 82.74 | 80.74 | 0.00 |
| age | 7 | 86.02 | 82.02 | 0.00 |
| $sex_{chick}$ | 7 | 86.18 | 82.18 | 0.00 |
| c | 6 | 89.24 | 83.23 | 0.00 |
| *(d) Distance from nest during contact* | | | | |
| **$sex_{parent}$ + age** | **7** | **0.01** | **0.00** | **0.73** |
| $sex_{chick}$ + $sex_{parent}$ + age | 8 | 0.00 | 1.99 | 0.27 |
| $sex_{parent}$ | 6 | 19.79 | 17.78 | 0.00 |
| $sex_{chick}$ + $sex_{parent}$ | 7 | 19.75 | 19.74 | 0.00 |
| age | 6 | 44.83 | 42.82 | 0.00 |
| $sex_{chick}$ + age | 7 | 44.82 | 44.81 | 0.00 |
| c | 5 | 64.56 | 60.55 | 0.00 |
| $sex_{chick}$ | 6 | 64.53 | 62.52 | 0.00 |

**Table S9.**

Model selection results with chick-parentID as levels for the random effects. The most parsimonious models are indicated in bold. "age" refers to age of the chick and "c" to the intercept-only "constant" model. For modelling contact probabilities, year was always included as a categorical fixed effect.

| Model | K | Δ-2logL | ΔAIC$_c$ | Akaike weight |
|---|---|---|---|---|
| *(a) Total contact* | | | | |
| sex$_{chick}$ + sex$_{parent}$ + age | 9 | 0.00 | 0.00 | 0.32 |
| sex$_{parent}$ + age | 8 | 2.36 | 0.35 | 0.27 |
| sex$_{chick}$ + age | 8 | 2.67 | 0.67 | 0.23 |
| **age** | **7** | **5.12** | **1.12** | **0.18** |
| sex$_{chick}$ + sex$_{parent}$ | 8 | 14.06 | 12.06 | 0.00 |
| sex$_{parent}$ | 7 | 16.39 | 12.39 | 0.00 |
| sex$_{chick}$ | 7 | 16.77 | 12.77 | 0.00 |
| c | 6 | 19.19 | 13.19 | 0.00 |
| *(b) Contact while begging* | | | | |
| **sex$_{chick}$ + age** | **8** | **1.32** | **0.00** | **0.40** |
| sex$_{chick}$ + sex$_{parent}$ + age | 9 | 0.00 | 0.68 | 0.28 |
| sex$_{parent}$ + age | 8 | 2.73 | 1.41 | 0.20 |
| age | 7 | 5.63 | 2.31 | 0.12 |
| sex$_{chick}$ | 7 | 17.27 | 13.95 | 0.00 |
| sex$_{chick}$ + sex$_{parent}$ | 8 | 15.81 | 14.49 | 0.00 |
| sex$_{parent}$ | 7 | 18.20 | 14.88 | 0.00 |
| c | 6 | 21.33 | 16.01 | 0.00 |
| *(c) Contact while foraging* | | | | |
| **age** | **7** | **0.05** | **0.00** | **0.50** |
| sex$_{parent}$ + age | 8 | 0.00 | 1.95 | 0.19 |
| sex$_{chick}$ + age | 8 | 0.05 | 2.00 | 0.19 |
| sex$_{chick}$ + sex$_{parent}$ + age | 9 | 0.00 | 3.95 | 0.07 |
| c | 6 | 8.01 | 5.95 | 0.03 |
| sex$_{parent}$ | 7 | 7.86 | 7.80 | 0.01 |
| sex$_{chick}$ | 7 | 7.90 | 7.85 | 0.01 |
| sex$_{chick}$ + sex$_{parent}$ | 8 | 7.74 | 9.69 | 0.00 |
| *(d) Distance from nest during contact* | | | | |
| **age** | **6** | **0.12** | **0.00** | **0.53** |
| sex$_{parent}$ + age | 7 | 0.00 | 1.88 | 0.21 |
| sex$_{chick}$ + age | 7 | 0.12 | 2.00 | 0.19 |
| sex$_{chick}$ + sex$_{parent}$ + age | 8 | 0.00 | 3.88 | 0.08 |
| c | 5 | 30.88 | 28.75 | 0.00 |
| sex$_{parent}$ | 6 | 30.82 | 30.70 | 0.00 |
| sex$_{chick}$ | 6 | 30.88 | 30.75 | 0.00 |
| sex$_{chick}$ + sex$_{parent}$ | 7 | 30.82 | 32.70 | 0.00 |

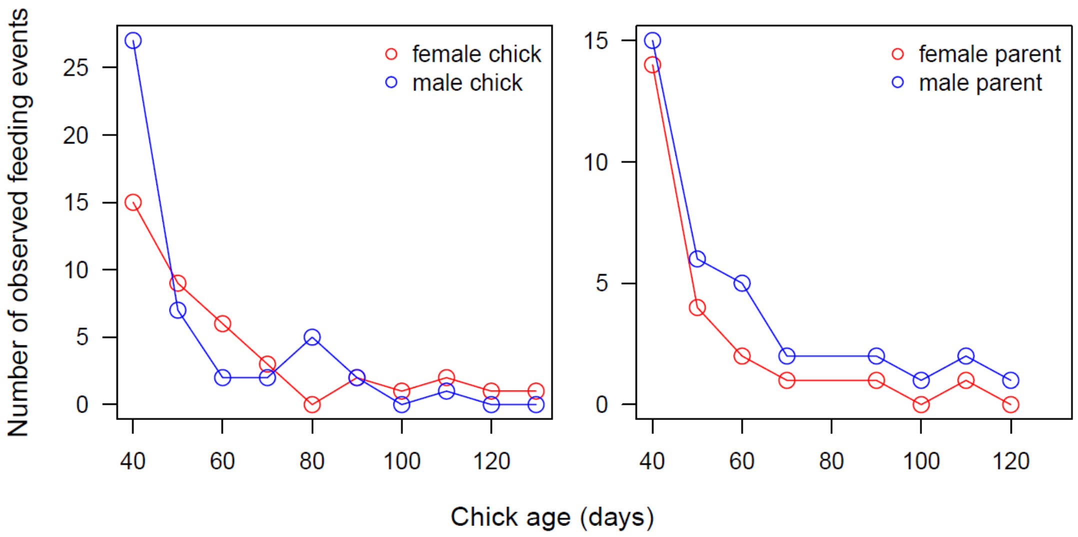

**Figure S1.**

Number of observed feeding events in relation to sex of the chick and the parent.

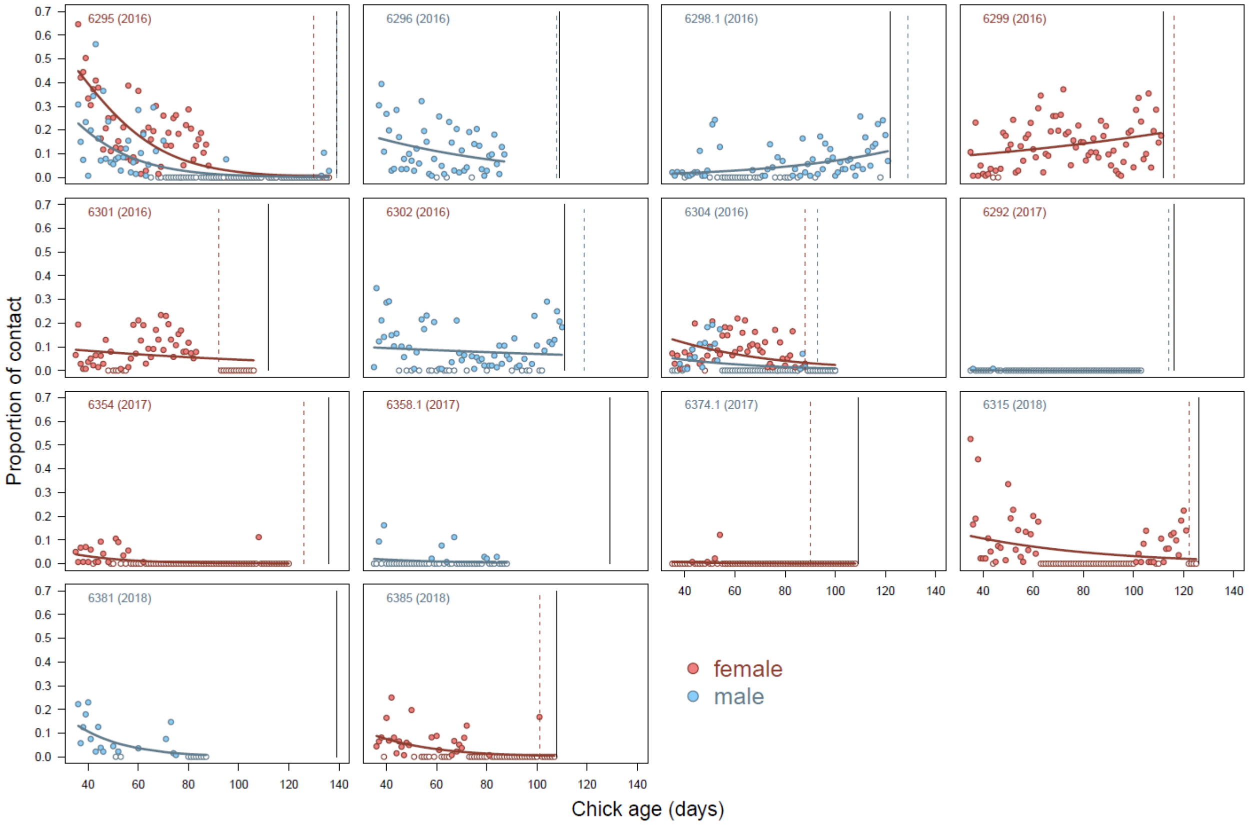

**Figure S2.**

The probability of contact between chick and parent in relation to chick age and coloured according to the chick's (chick number's color) and parent's sex (symbol color). Solid colored lines represent model estimates from the most parsimonious model, the vertical lines represent the moments of departure of the chick (black line) and the parent (colored dashed line). Days without any contact are plotted as white-filled circles.

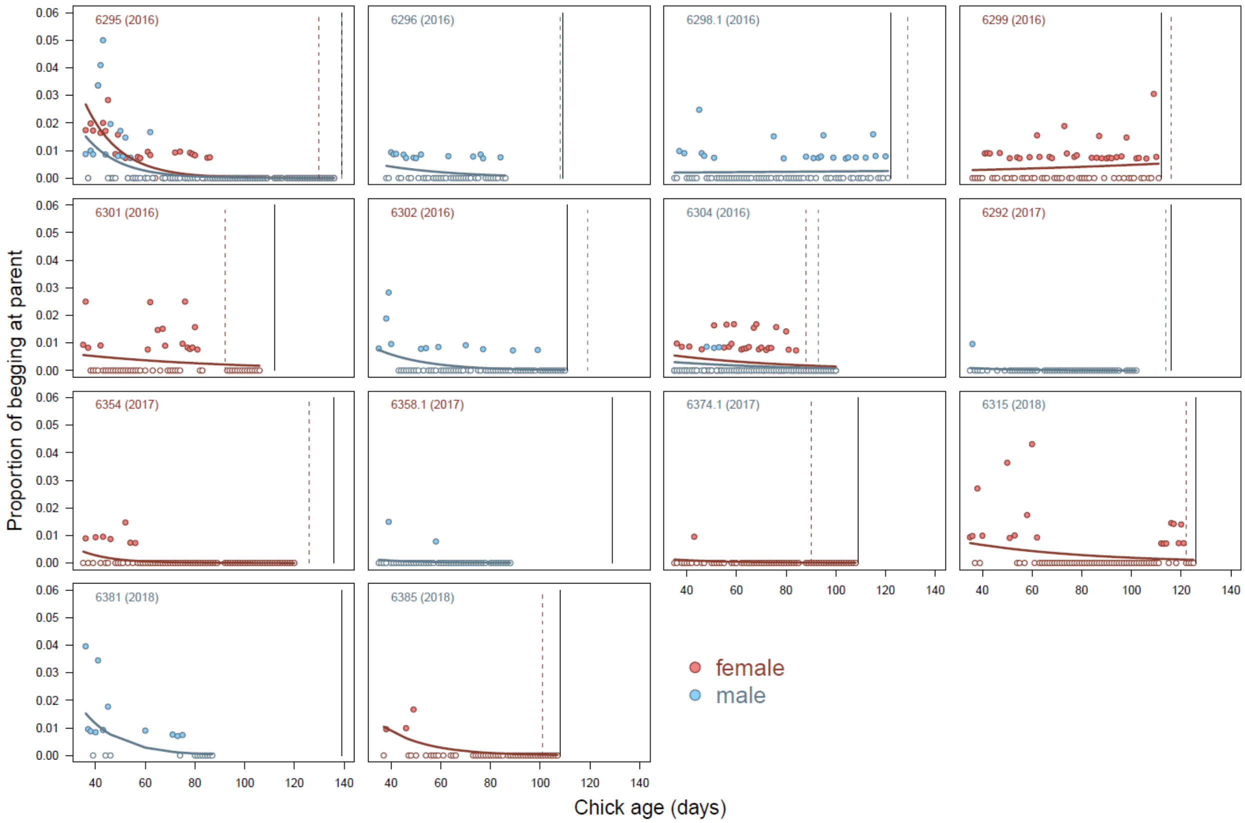

**Figure S3.**

The probability of contact between chick and parent while the chick is classified to be begging in relation to chick age and coloured according to the chick's (chick number's color) and parent's sex (symbol color). Solid colored lines represent model estimates from the most parsimonious model, the vertical lines represent the moments of departure of the chick (black line) and the parent (colored dashed line). Days without any 'begging contact' are plotted as white-filled circles.

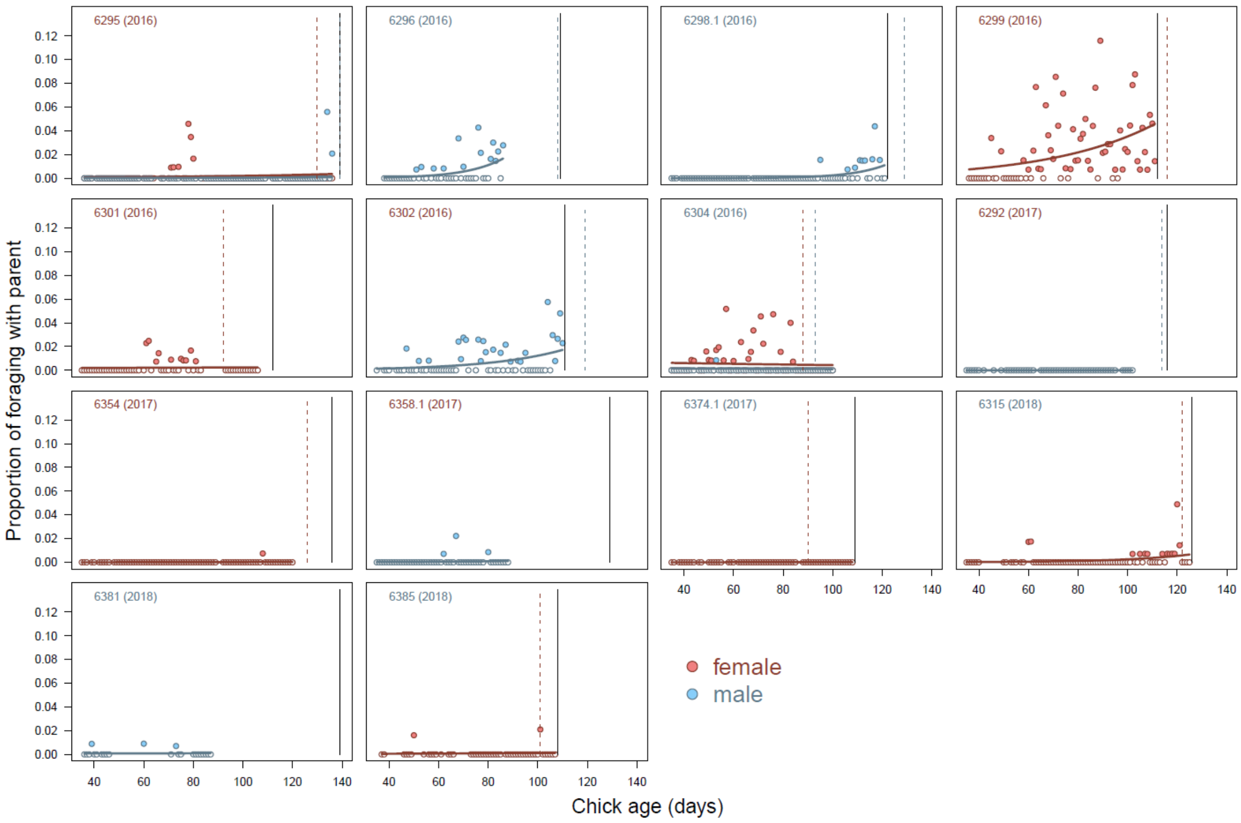

**Figure S4.**

The probability of contact between chick and parent while the chick is classified to be foraging in relation to chick age and coloured according to the chick's (chick number's color) and parent's sex (symbol color). Solid colored lines represent model estimates from the most parsimonious model, the vertical lines represent the moments of departure of the chick (black line) and the parent (colored dashed line). Days without any 'foraging' contact are plotted as white-filled circles.
